# Codon clusters with biased synonymous codon usage represent hidden functional domains in protein-coding DNA sequences

**DOI:** 10.1101/530345

**Authors:** Zhen Peng, Yehuda Ben-Shahar

## Abstract

Protein-coding DNA sequences are thought to primarily affect phenotypes via the peptides they encode. Yet, emerging data suggest that, although they do not affect protein sequences, synonymous mutations can cause phenotypic changes. Previously, we have shown that signatures of selection on gene-specific codons usage bias are common in genomes of diverse eukaryotic species. Thus, synonymous codon usage, just as amino acid usage pattern, is likely a regular target of natural selection. Consequently, here we propose the hypothesis that at least for some protein-coding genes, codon clusters with biased synonymous codon usage patterns might represent “hidden” nucleic-acid-level functional domains that affect the action of the corresponding proteins via diverse hypothetical mechanisms. To test our hypothesis, we used computational approaches to identify over 3,000 putatively functional codon clusters (PFCCs) with biased usage patterns in about 1,500 protein-coding genes in the *Drosophila melanogaster* genome. Specifically, our data suggest that these PFCCs are likely associated with specific categories of gene function, including enrichment in genes that encode membrane-bound and secreted proteins. Yet, the majority of the PFCCs that we have identified are not associated with previously annotated functional protein domains. Although the specific functional significance of the majority of the PFCCs we have identified remains unknown, we show that in the highly conserved family of voltage-gated sodium channels, the existence of rare-codon cluster(s) in the nucleic-acid region that encodes the cytoplasmic loop that constitutes inactivation gate is conserved across paralogs as well as orthologs across distant animal species. Together, our findings suggest that codon clusters with biased usage patterns likely represent “hidden” nucleic-acid-level functional domains that cannot be simply predicted from the amino acid sequences they encode. Therefore, it is likely that on the evolutionary timescale, protein-coding DNA sequences are shaped by both amino-acid-dependent and codon-usage-dependent selective forces.

## 2. Introduction

In general, it is assumed that the primary function of a protein-coding sequence is to encode a specific sequence of amino acids whose biochemical properties determine the structure and functions of the encoded peptide. However, emerging data indicate that synonymous mutations, which do not affect amino acid sequences, can still have dramatic phenotypic impacts. Thus, it has been hypothesized that some important factors affecting protein structures and functions are not simply encoded by amino acid residues but by nucleic-acid-level information, such as codon usage bias [1,2]. Therefore, just as a sequence of amino acids with a specific order and/or specific biochemical properties can form a protein domain that performs specific functions, it is also possible that a sequence of codons with a specific codon usage pattern could serve as a nucleic-acid-level domain that affects the functions of the mature protein.

Based on the hypothesis that codon-usage-encoded domains can affect protein functions, researchers have identified rare-codon clusters, defined by whole-genome codon usage frequencies, that possibly decelerate translation and thus modify protein functions by affecting co-translational folding and/or modifications of nascent peptide chains [2–7]. Nevertheless, if functional codon clusters do exist, local deceleration of translation may not be the only mechanism through which they affect protein functions. It is also possible that functional codon clusters could correspond to locally accelerated translation, a specific combination of translationally decelerated and accelerated regions, specific RNA secondary structures [8,9], and binding sites for miRNAs [10]. Thus, for generally investigating codon clusters as functional domains that may be “hidden” from the amino acid sequences, exclusive focus on rare-codon clusters may lead to biased results. Therefore, it is necessary to develop statistical methods that generally detect putatively functional codon clusters (PFCCs), no matter what specific codons they prefer or through what mechanisms they may affect protein functions.

Consequently, to identify PFCCs, we developed a conservative statistical approach and applied it to the *Drosophila melanogaster* genome with approximately 14,000 protein-coding genes, which yielded over 3,000 PFCCs in about 1,500 genes. Interestingly, some of these PFCCs strongly prefer common codons while some others adopt complex codon usage patterns that cannot be simply described as preference for common or rare codons, which has not been reported before. Furthermore, we found that genes encoding transmembrane proteins are more likely to bear PFCCs. However, only a small proportion of the identified PFCCs are associated with the coding sequences of transmembrane helices, which suggests that PFCCs are either associated with other types of protein domains that are overrepresented in transmembrane proteins or not necessarily associated with amino-acid-encoded domains. We further found that the majority of the identified PFCCs are not associated with established protein domains in the Pfam database [11]. These data suggest that most PFCCs likely encode “hidden” nucleic-acid-level functional domains that cannot be predicted solely from amino acid sequences. The rationale for this inference is as follows: first, Pfam is a well-established database of conserved protein domains that have undergone strong natural selection; second, the PFCCs can be identified only when natural selection on local codon usage patterns is strong enough to generate statistically detectable signals; third, if the major impacts of PFCCs on gene functions are mediated by amino-acid-encoded protein domains, most PFCCs are expected to be associated with amino-acid-encoded domains that have undergone strong natural selection; fourth, the actual observation contradicts the expectation, and thus the functions of PFCCs should not be strongly associated with amino-acid-encoded domains. Finally, by implementing comparative analysis between homologs, we showed that the family of voltage-gated sodium channels likely evolved conserved preference for rare codons in a region responsible for the channel inactivation. Together, our data suggest that similar to amino acid sequences, codon clusters can also encode diverse functional domains, which provides an additional level of regulation over the structures, modifications, and functions of proteins.

## 3. Results

### 3.1. Identifying putatively functional codon clusters (PFCCs)

If the synonymous codon usage of a codon cluster does not perform specific functions, it should not be affected by natural selection and thus it can be explained by the background codon usage frequencies, which is mainly determined by mutations and genetic drift [12,13]. For example, if a gene locates in a GC-enriched chromosomal region that has resulted from GC-biased mutations, it is expected that the background codon usage are biased towards GC-ended codons; thus, if a sub-genic region is not significantly affected by natural selection on codon usage, its synonymous codon usage should also be biased towards GC-ended codons. Therefore, if the codon usage pattern of a codon cluster cannot be explained by the background codon usage frequencies, it should be significantly affected by natural selection; thus, such a codon cluster is by definition a PFCC. To identify PFCCs, first we needed to choose background codon usage frequencies. Previous studies on synonymous codon usage usually used the whole-genome codon usage frequencies as the background [3,5–7]. Nevertheless, our recent study [14] showed that gene-specific codon usage pattern can be fairly different from whole-genome one. Thus, a codon cluster whose synonymous codon usage cannot be explained by whole-genome codon usage may be adequately explained by gene-specific codon usage, and *vice versa*. Therefore, to filter out the interference from the discrepancies between whole-genome and gene-specific codon usage patterns so that PFCCs are conservatively identified, neither whole-genome nor gene-specific codon usage frequencies should be able to explain the codon usage pattern of a PFCC. Based on the aforementioned logic, we developed a statistical approach to scan protein-coding sequences in order to identify PFCCs (see Materials and Methods: Identifying PFCCs).

By applying the approach to 13,821 protein-coding genes from the reference *D. melanogaster* genome, we identified 3,050 PFCCs in 1,445 genes (Table S1). This result indicates that PFCCs do exist, and they impact at least 10% of protein-coding genes in the *D. melanogaster* genome.

### 3.2. Codon usage patterns of PFCCs are diverse

In principle, the codon usage patterns of PFCCs can deviate from the background codon usage frequencies for various non-mutually exclusive biological reasons. First, the enrichment of rare codons in a PFCC might decelerate translation [7]. Second, it is possible that the enrichment of common codons in a PFCC could accelerate translation. Third, PFCCs with more complex codon usage patterns, which cannot be simply described as the preference for common or rare codons, might serve important functions by modifying mRNA secondary structure [8,9] or miRNA accessibility [10]. Thus, classifying the identified PFCCs by their codon usage patterns could be informative for assessing how PFCCs may affect protein functions.

Codon adaptation index (CAI) [15] has been widely used to describe a protein-coding sequence’s propensity of using common codons. In general, a higher CAI indicates stronger preference for common codons and/or avoidance of rare codons. However, directly using CAI as the index to classify PFCCs could lead to biased results, especially when common codons are not enriched in the PFCCs. This is because the differences between usage frequencies of the synonymous codons for some amino acids are much larger than those of other amino acids. Thus, even if two codon clusters both strictly use rare codons, they could have very different CAIs depending on the amino acid sequences. To circumvent such a weakness of CAI, we propose to use a transformed CAI (TCAI) to describe the general codon usage pattern of a PFCC.

TCAI is calculated as below. For a PFCC, the corresponding amino acid sequence and the background codon usage pattern – either the whole-genome or gene-specific codon usage pattern – are used to randomly generate 10,000 “pseudo-clusters” of codons that encode exactly the same amino acid sequences as what is encoded by the PFCC. Thus, on average, the overall codon usage patterns of these pseudo-clusters should follow the background codon usage pattern. Then the CAIs of all pseudo-clusters are calculated, and TCAI is defined as the result of subtracting the proportion of pseudo-clusters whose CAIs are higher than the CAI of the PFCC from the proportion of pseudo-clusters whose CAIs are lower than the CAI of the PFCC. Thus, TCAI varies between −1 and 1. TCAI = −1 means that the CAIs of all pseudo-clusters are higher than that of the PFCC, suggesting that the PFCC strongly prefers rare codons; in contrast, TCAI =1 suggests that the PFCC strongly prefers common codons. Thus, TCAI effectively suppresses the interference from different levels of codon usage biases for different amino acids.

We calculated TCAIs for all identified PFCCs, either by using whole-genome (Fig. 1A) or gene-specific (Fig. 1B) codon usage pattern as the background. The distribution of TCAI values (Fig. 1) indicates that most of the PFCCs are rare-codon clusters, while common-codon clusters do exist as shown by a small peak in the rightmost part of the histograms. More interestingly, there are also some codon clusters whose TCAI values are intermediate, suggesting that their codon usage patterns are more complex and cannot be simply described by strong preference for common or rare codons. The preponderance of rare-codon clusters may be explained by two reasons that are not mutually exclusive. First, the preponderance may represent the fact that rare-codon clusters are biologically more important than other types of functional codon clusters. Second, the preponderance may also be partly an artifact of technically easier detection of enriched rare codons in a short nucleotide sequence. Nonetheless, it was undoubtedly confirmed that there are different types of codon clusters in terms of synonymous codon usage patterns.

**Fig. 1.**
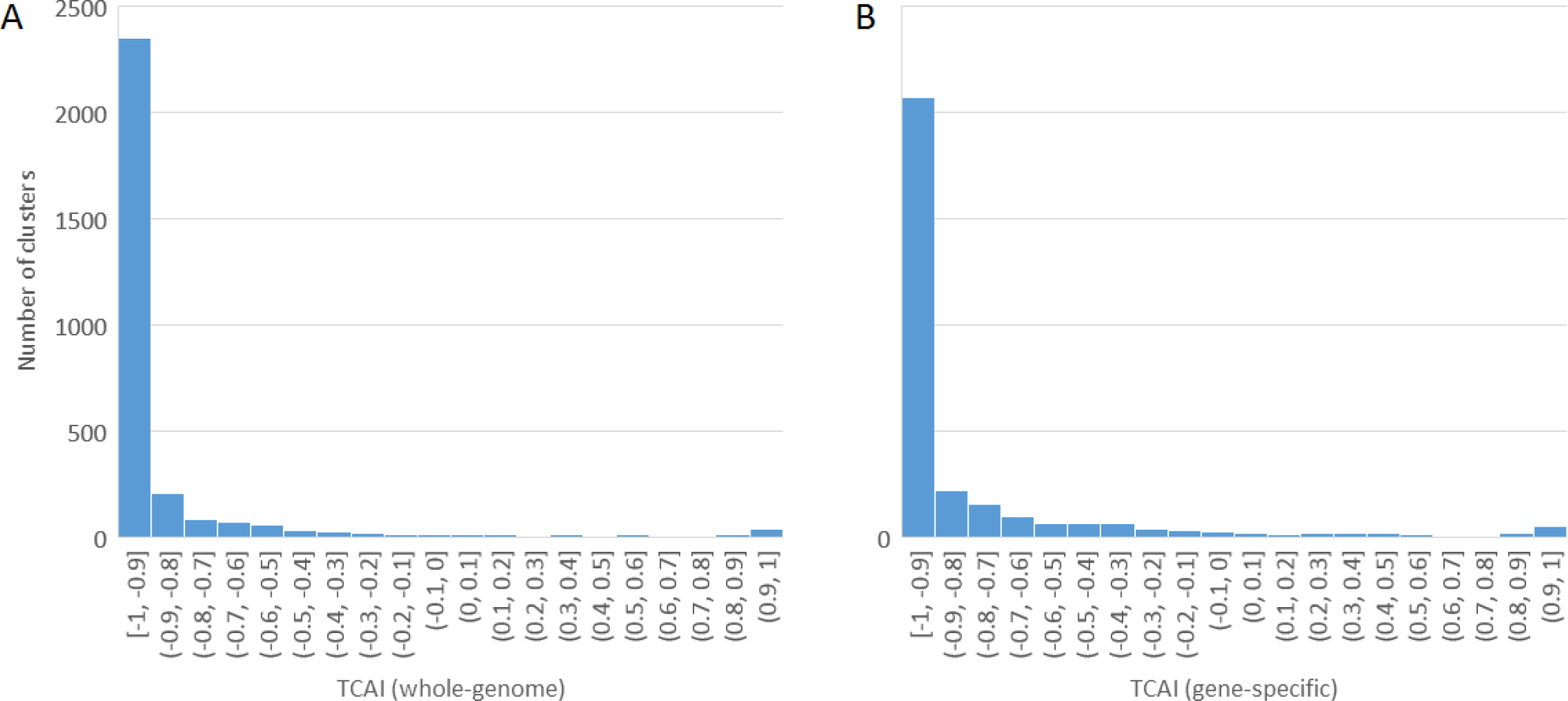
Distribution of TCAI values. TCAI values were calculated by using the whole-genome (**A**) or gene-specific (**B**) codon usage patterns as the background codon usage. The TCAI of a rare-codon cluster is near −1, while that of a common-codon cluster is near 1.

We also noted that although the distribution patterns shown in Fig. 1A and Fig. 1B are qualitatively similar, the actual values of corresponding columns in the histograms are quantitatively different. This suggested the possibility that a PFCC could be assigned to different types of codon clusters, depending on which background codon usage pattern was used. Such a possibility may interfere the interpretations of the putative functions of the PFCC. For example, a rare-codon cluster in terms of whole-genome codon usage may be classified as a common-codon cluster in terms of gene-specific codon usage, and thus it could be unclear whether the PFCC may decelerate or accelerate translation. In order to assess the influence of the discrepancy between whole-genome and gene-specific codon usage patterns on the classification of PFCCs, we used a scatter plot to examine the relationship between whole-genome TCAI and gene-specific TCAI (Fig. 2). The data points were then clustered by K-mean clustering to seven types (K=7).

**Fig. 2.**
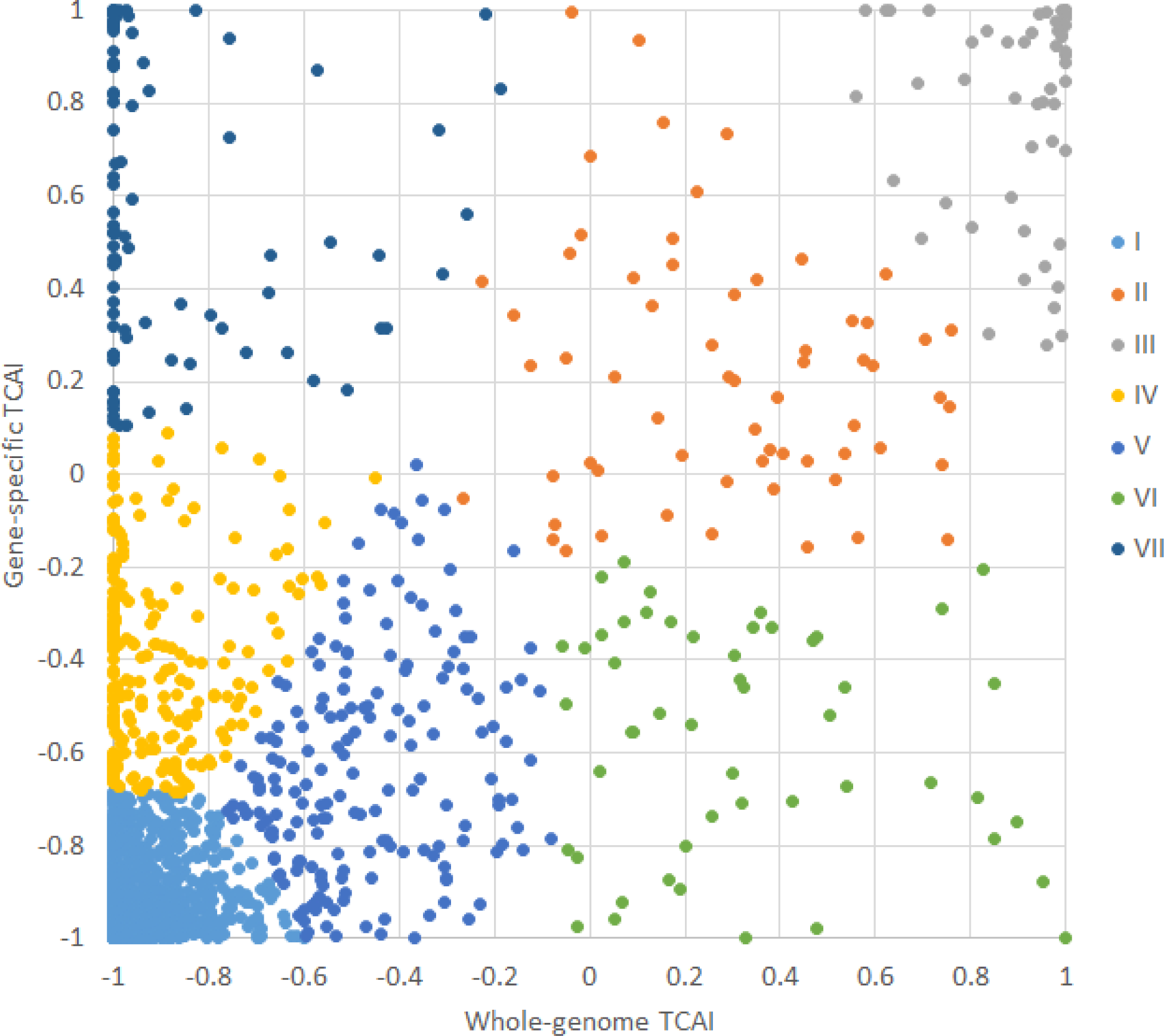
Influence of the discrepancy between whole-genome and gene-specific codon usage patterns on classifying PFCCs. Codon usage patterns of identified PFCCs were described by TCAI. Since gene-specific and whole-genome-level TCAI values for the same codon cluster could be different, we plotted the gene-specific TCAI against whole-genome TCAI for all codon clusters and then classified codon clusters by K-mean clustering (K=7).

We found that most codon clusters have similar whole-genome and gene-specific TCAI (Fig. 2, types I-V). However, some common-codon clusters in terms of whole-genome TCAI were classified as rare-codon clusters in terms of gene-specific TCAI (Fig. 2, type VI), and *vice versa* (Fig. 2, type VII). This result suggests that due to the discrepancy between whole-genome and gene-specific codon usage patterns, it is difficult to predict the exact biological roles of some identified PFCCs. For example, in our previous study, we showed that some whole-genome rare codons can be common and translationally optimal for tissue-specific genes [14]. Thus, a rare-codon cluster in terms of whole-genome codon usage, which would be naïvely considered as a “decelerating codon cluster”, might be a common-codon cluster in terms of gene-specific codon usage, which could actually serve as an “accelerating codon cluster”. Therefore, although PFCCs can be detected by statistical approaches proposed by us and others [6,7], to computationally predict the candidate functional roles of these codon clusters may require extra information such as tRNA expression profile and better tools for predicting the secondary and tertiary structures of RNAs.

To summarize, PFCCs are diverse according to their codon usage patterns. Rare-codon clusters, whose main function is presumably decelerating translation [6,7], seem to be the majority of PFCCs. There are also other types of PFCCs, including common-codon clusters and PFCCs with more complex codon usage patterns, which likely have functions other than decelerating translation. Nonetheless, the discrepancy between whole-genome and gene-specific codon usage patterns makes it hard to predict the possible functions of the PFCCs whose whole-genome TCAI and gene-specific TCAI are dramatically different.

### 3.3. PFCC distribution is not restricted to specific regions of protein-coding sequences

Except for the codon usage patterns of PFCCs, the locations of PFCCs in protein-coding sequences may also provide hints to the possible functions of PFCCs. Previous studies have shown that a potential important function of codon clusters is that N-terminal rare-codon clusters could affect secretion of proteins [16–18], possibly via interaction with the nascent chains of signal peptides [17,18]. Therefore, we next tested if the PFCCs detected by our approach tend to locate near the N-terminus; if they do, it could suggest that PFCCs are likely associated with secretion of proteins.

To measure how close a PFCC-encoded region is to the N-terminus, we defined the relative location index (RLI) of a PFCC as the ratio of the distance between the midpoint of the PFCC-encoded region and the N-terminus to the length of the entire protein. Thus, a small RLI means that the PFCC-encoded region is close to the N-terminus. We then plotted the distribution of PFCCs against their RLIs (Fig. 3A). We found that although the density of PFCCs is apparently higher in the N-terminal region, the distribution of PFCCs is not restricted to this region (Fig. 3A). As we have assigned these codon clusters to seven types (Fig. 2), we also examined if some specific types of PFCCs exhibit skewed distribution towards the N-terminal region (Fig. 3B-H). As expected, type I codon clusters, which can be described as rare-codon clusters, exhibit slight enrichment near the N-terminus (Fig. 3B). To our surprise, type III codon clusters, which can be described as common-codon clusters, also exhibit relatively strong enrichment near the N-terminus (Fig. 3D). Other types of codon clusters do not exhibit clear enrichment near the N-terminus. We also performed a gene ontology (GO) analysis [19,20] (http://geneontology.org/) on the genes carrying N-terminal codon clusters (RLI < 0.1) to see if the genes encoding secreted proteins are enriched. We found that not only some extracellular matrix structural constituents, mostly mucins, are enriched, but also proteins associated with plasma membrane or transcription-level regulation are enriched (Table S2).

**Fig. 3.**
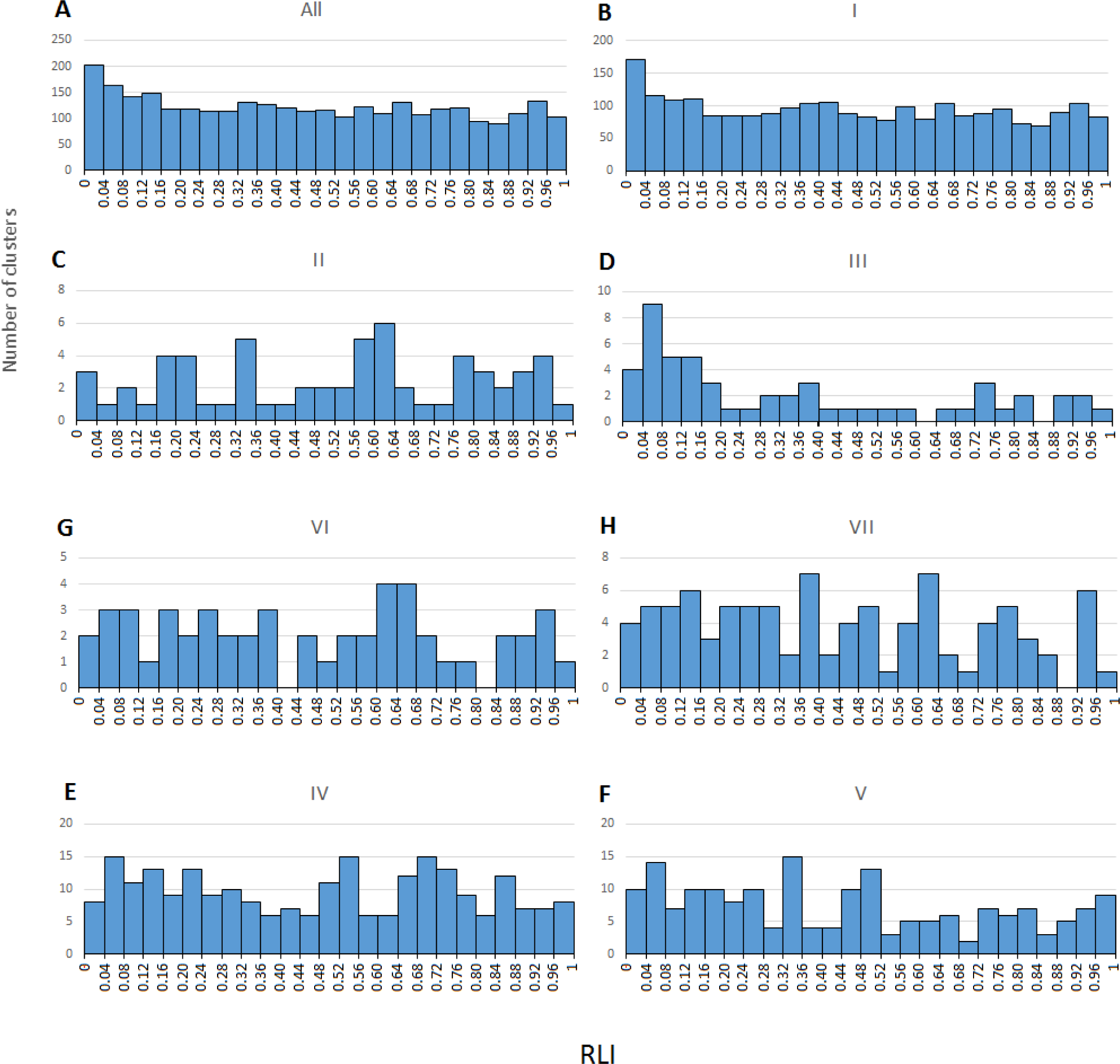
Spatial distribution of putatively functional codon clusters. For all identified PFCCs and each type of PFCCs shown in Fig. 2, the distribution of PFCCs is plotted against the location coordinates, measured by RLI (RLI=0 means N-terminus; RLI=1 means C-terminus).

Together, these data indicate that although N-terminal regions are more likely to harbor PFCCs, many PFCCs actually locate in other regions (Fig. 3). They also suggest that although the function of a subset of the PFCCs may be explained by N-terminal rare-codon clusters’ impact on secretion or signal peptides, such a function is unlikely a general role played by other PFCCs. For example, the codon clusters locating in the middle of genes should have little to do with signal peptides. Thus, PFCCs likely perform various biological functions that need further investigation.

### 3.4. Specific protein functional classes are overrepresented in genes carrying PFCCs while most PFCCs are not associated with known protein domains

To further investigate the biological roles of PFCCs, we next performed GO analyses on the genes carrying PFCCs, in order to test the hypothesis that PFCCs are associated with various functional features of protein-coding genes. We found that in all 1445 genes that carry the PFCCs, genes encoding membrane-binding proteins and transcription-related proteins are overrepresented, while genes encoding ribosomal and mitochondrial proteins are underrepresented (Table S3). This result suggests that functional codon clusters might be associated with transmembrane domains, so we then tested if the amino acid sequences encoded by the PFCCs are near or overlapped with the transmembrane helices predicted by TMHMM, an algorithm predicting transmembrane helices [21]. Unexpectedly, we found that only about 6% of the PFCCs are near or overlapped with some transmembrane helices (Table 1). Thus, there seems to be a discrepancy between the overrepresentation of transmembrane proteins in the genes carrying PFCCs and relatively few PFCCs that are near or overlapped with the sequences encoding transmembrane helices. Nevertheless, such a discrepancy could be explained by that PFCCs may be functionally more important for the non-transmembrane regions in transmembrane proteins. The discrepancy may also be explained by that transmembrane helices are less sensitive to the change in codon usage since the helices are strongly affected by the biochemical properties, such as hydrophobicity, of amino acid residues [21,22].

**Table 1.**
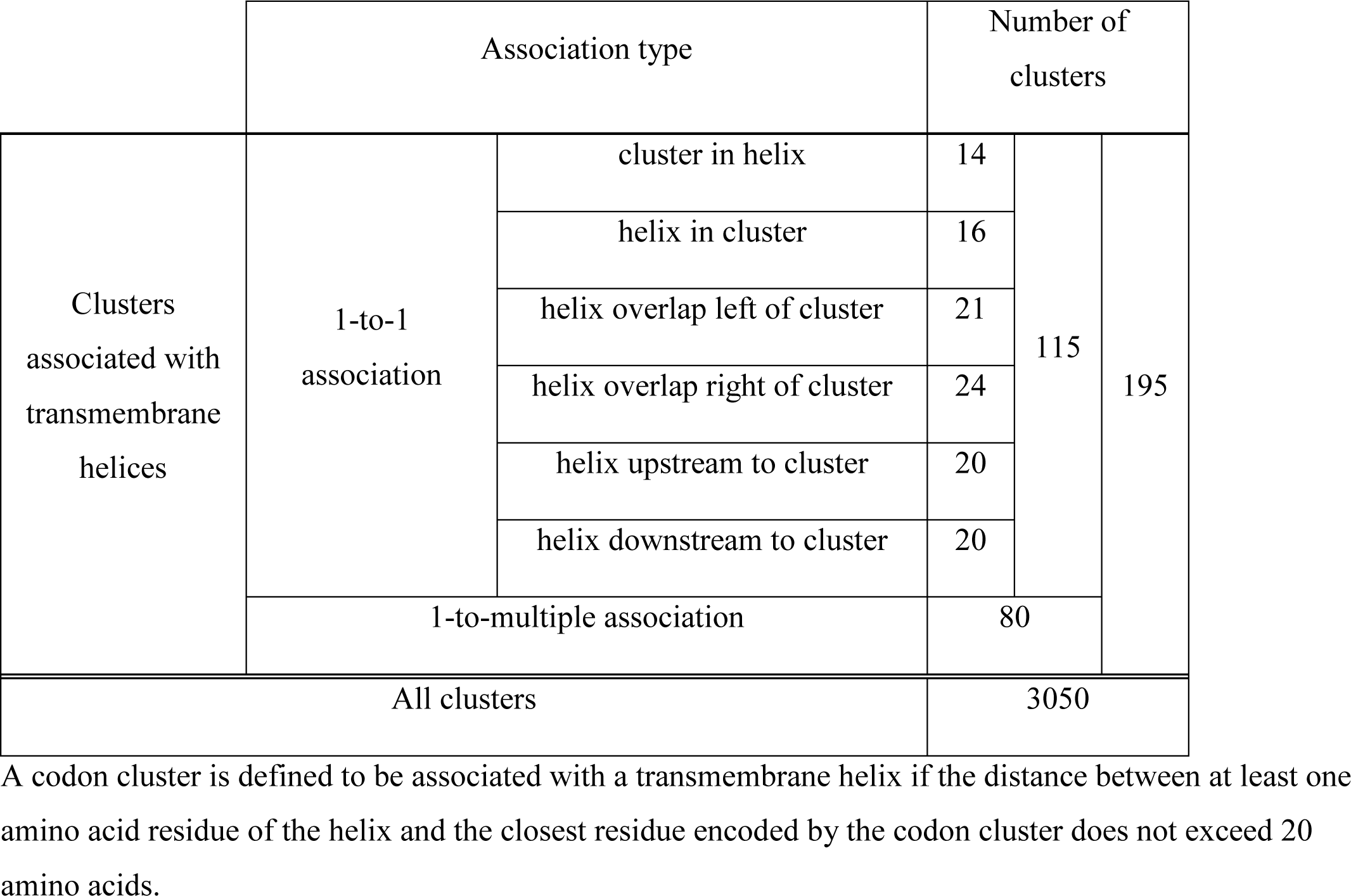
**Biased codon clusters overlap with transmembrane helices.**

|  | Association type |  | Number of clusters |  |  |
| --- | --- | --- | --- | --- | --- |
| Clusters associated with transmembrane helices | 1-to-1 association | cluster in helix | 14 | 115 | 195 |
|  |  | helix in cluster | 16 |  |  |
|  |  | helix overlap left of cluster | 21 |  |  |
|  |  | helix overlap right of cluster | 24 |  |  |
|  |  | helix upstream to cluster | 20 |  |  |
|  |  | helix downstream to cluster | 20 |  |  |
|  | 1-to-multiple association |  | 80 |  |  |
| All clusters |  |  | 3050 |  |  |

A codon cluster is defined to be associated with a transmembrane helix if the distance between at least one amino acid residue of the helix and the closest residue encoded by the codon cluster does not exceed 20 amino acids.

If most PFCCs are not associated with transmembrane helices, then what other types of protein domains might be associated with the PFCCs? To answer this question, we examined the association between PFCCs and annotated protein domains in the Pfam database [11,23]. We found that about 1/4 of the PFCCs are near or overlapped with some annotated Pfam protein domains, yet it is still unclear how the other 3/4 might influence protein functions (Table 2, Table S4). Among the PFCCs of which each is associated with only one Pfam protein domain, about 1/2 locate within protein domains (Table 2), which was consistent with what was recently reported by Chaney et al. [7]. These data suggest that although some PFCCs likely affect protein functions by modifying the co-translational processes concerning protein domains defined by amino acid sequences, the majority of PFCCs seem to be associated with unknown functional domains.

**Table 2.** **Biased codon clusters overlap with Pfam domains.**

|  | Association type |  | Number of clusters |  |  |
| --- | --- | --- | --- | --- | --- |
| Clusters associated with Pfam domains | 1-to-1 association | cluster in domain | 299 | 584 | 746 |
|  |  | domain in cluster | 3 |  |  |
|  |  | domain overlap left of cluster | 63 |  |  |
|  |  | domain overlap right of cluster | 75 |  |  |
|  |  | domain upstream to cluster | 58 |  |  |
|  |  | domain downstream to cluster | 86 |  |  |
|  | 1-to-multiple association |  | 162 |  |  |
| All clusters |  |  | 3050 |  |  |

A codon cluster is defined to be associated with a Pfam domain if the distance between at least one amino acid residue of the Pfam domain and the closest residue encoded by the codon cluster does not exceed 20 amino acids.

To summarize, although specific protein functional classes are overrepresented in the genes carrying PFCCs, most of the PFCCs are not associated with known protein domains defined by amino acid sequences. Therefore, PFCCs likely represent “hidden” nucleic-acid-level domains that regulate protein functions.

### 3.5. Voltage-gated sodium channels include a conserved rare-codon cluster associated with the inactivation gate

To identify possible specific functions of some PFCCs, we next investigated PFCCs identified in the *D. melanogaster* voltage-gated sodium channel (Nav) genes as a proof of principle for the following reasons. First, Nav has multiple transmembrane domains [24–26] and we have shown that transmembrane proteins are associated with PFCCs (Table S3). Second, Nav is a well-characterized protein family in terms of its physiological roles and structure-function relationship. Third, the *D. melanogaster* genome harbors two Nav paralogs whose divergence was dated back to the origin of Bilateria, which allows us to identify the PFCCs with conserved codon usage patterns.

Each Nav α-subunit consists of four transmembrane domains (Domains I-IV) linked by cytoplasmic chains, plus an N-terminal and a C-terminal cytoplasmic chains. The inactivation gate, which is responsible for stopping the sodium influx during action potential, is formed by the cytoplasmic chain between Domain III (DIII) and Domain IV (DIV) that will be refer to as DIII-IV linker below [26]. In general, most invertebrates have two types of Nav, namely type 1 Nav (Nav1) and type 2 Nav (Nav2), while vertebrates have lost the Nav2 gene but have gained multiple Nav1 paralogs [26]. As aforementioned, *D. melanogaster* has two paralogs of Nav, namely *para*, the Dmel/Nav1, and *NaCP60E*, the Dmel/Nav2 [27,28].

Multiple PFCCs were identified in Dmel/Nav1 and Dmel/Nav2, but the PFCCs in Dmel/Nav1 and those in Dmel/Nav2 are not always homologous. Nonetheless, we found that both genes have PFCCs in the DIII-IV linkers (Fig. 4). To assess the potential functions of these PFCCs, we then scanned the DIII-IV linkers with a 15-amino-acid sliding window and calculated TCAI for each window. We found that these PFCCs exhibit strong preference for rare codons (Fig. 5A, Dmel; Fig. 5B, Dmel), suggesting that decelerating translation during synthesizing the inactivation gate may be the key function of these PFCCs. We further scanned the DIII-IV linkers of Nav homologs in several other representative eukaryotic species, and found that the majority of them also have sub-regions preferring rare codons (Fig. 5, TCAI < −0.8).

**Fig. 4.**
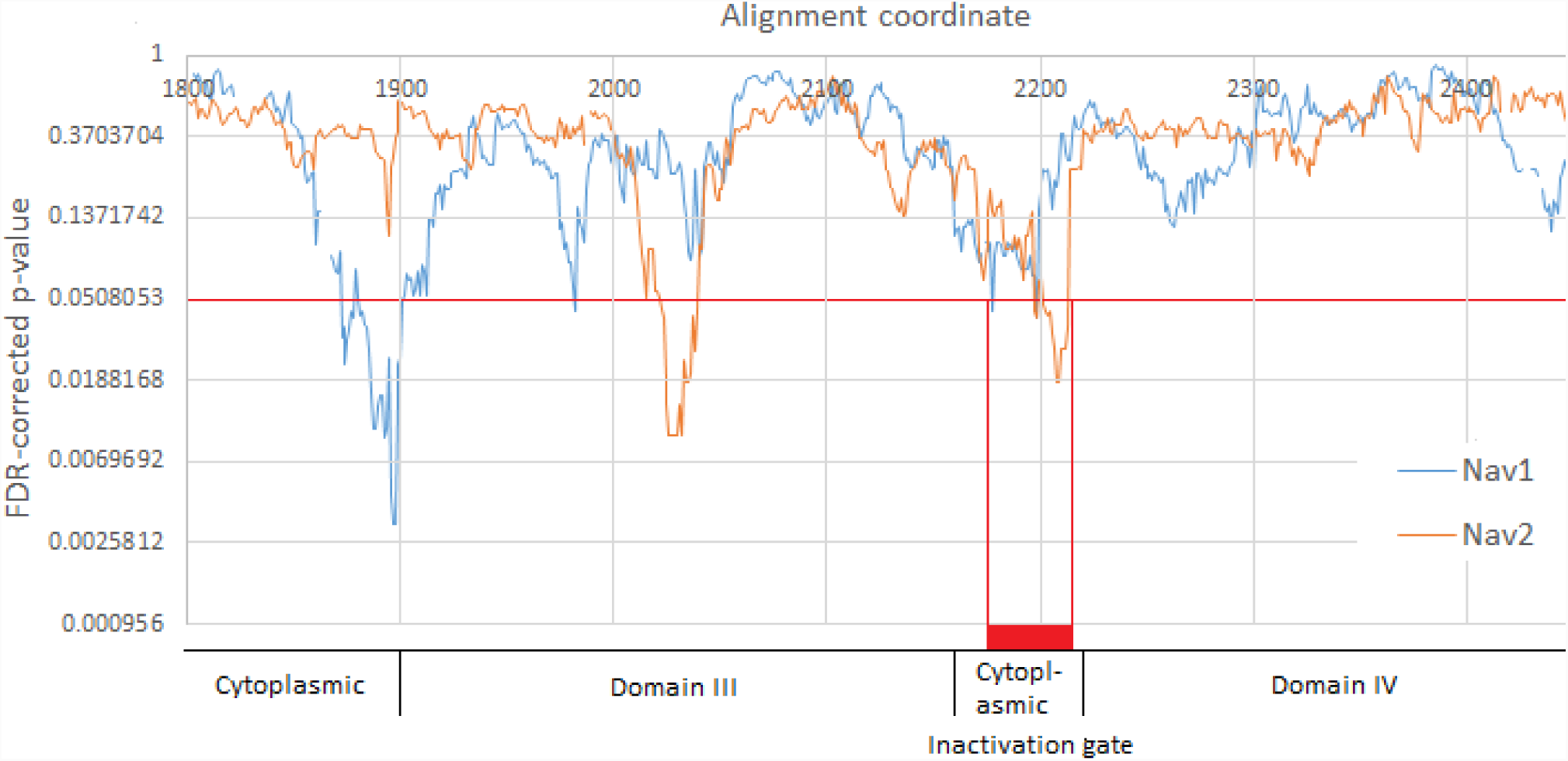
Identifying PFCCs in *D. melanogaster* Nav paralogs. Dmel/Nav1 and Dmel/Nav2 are aligned by amino acid sequences. *p*-values were corrected by FDR (FDR = 0.05), and those lower than the threshold indicate codon clusters whose codon usage patterns are significantly different from both whole-genome and gene-specific codon usage patterns. Both Dmel/Nav1 and Dmel/Nav2 have PFCCs in the DIII-IV linkers, shown by the red bar.

**Fig. 5.**
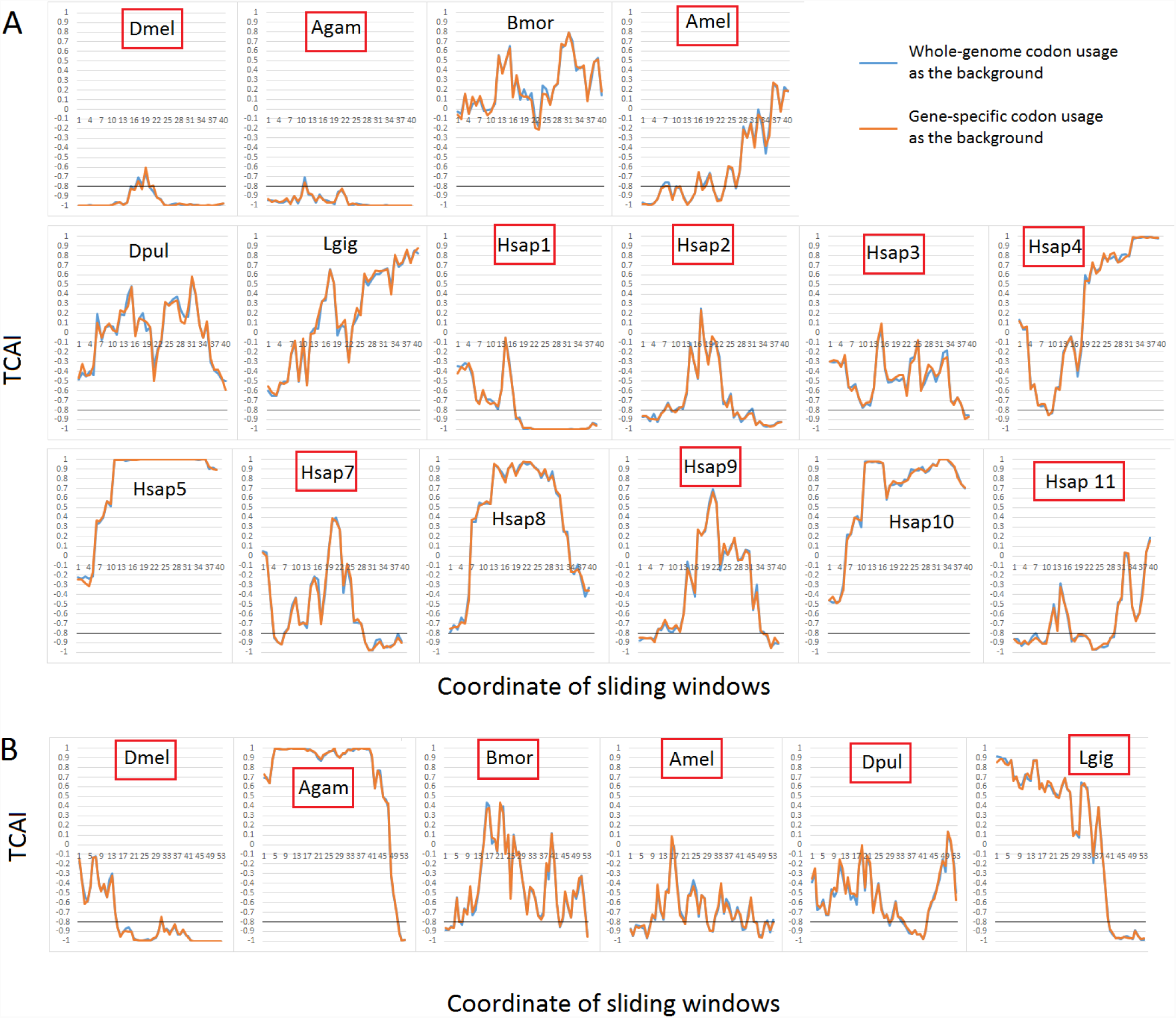
Nav paralogs generally bear rare-codon clusters in DIII-DIV linkers. **(A)** Nav1. **(B)** Nav2. Dmel: *Drosophila melanogaster*, fruit fly; Agam: *Anopheles gambiae*, malaria mosquito; Bmor: *Bombyx mori*, silkmoth; Amel: *Apis mellifera*, Western honey bee; Dpul: *Daphnia pulex*, water flea; Lgig: *Lottia gigantea*, owl limpet; Hsap: *Homo sapiens*, human. Homo sapiens has ten Nav1 paralogs but no Nav2. As suggested by Fig. 2, regions with TCAI < −0.8 are regarded as rare-codon clusters. Red boxes highlight the DIII-IV linkers carrying rare-codon clusters. Black lines: TCAI = 0.8. Blue curves: TCAI calculated by using whole-genome codon usage as the background. Orange curves: TCAI calculated by using gene-specific codon usage as the background.

Considering that the divergence between Nav1 and Nav2 was dated back to the origin of Bilateria [26], the conserved preference for rare codons in the DIII-IV linkers further support the hypothesis that the normal function of inactivation gate requires decelerated translation of this region. Decelerated translation is possibly critical for the correct folding pattern or phosphorylation of the DIII-IV linker [29–32]. In this regard, we hypothesize that synonymous mutations from rare codons to common codons in the DIII-IV linker could induce changes in the action potential through prolonged or shortened depolarization. Also, as some nonsynonymous mutations in the DIII-IV linker could cause cold-induced paralysis [33], it is possible that the synonymous mutations from rare codons to common codons in this region can cause similar phenotypes.

Furthermore, we noticed that not all DIII-IV linkers bear obvious rare-codon clusters (Fig. 5A, Bmor, Dpul, Lgig, Hsap5, Hsap8, Hsap10). Therefore, it is possible that for some species, synonymous codon usage in the DIII-IV linker is less sensitive to natural selection, perhaps due to other mechanisms that compensate the effects of rare codons on protein folding. More interestingly, we found that among the Nav1 paralogs in human, some have rare-codon clusters in the DIII-IV linkers while others do not. We also found that among the paralogs with rare-codon clusters, the specific locations of rare-codon clusters can be different. These findings perhaps suggest that rare-codon clusters are associated with the division of labor between Nav1 paralogs. As Nav1 paralogs have differentiated tissue-specific expression profiles [34], one mechanism underlying the possible codon-usage-mediated division of labor may be that these paralogs adapt their DIII-IV linkers’ codon usage patterns to tissue-specific tRNA pools [14,35,36], so that the corresponding protein-coding sequences are able to more finely regulate the function of inactivation gate.

## 4. Discussion

Here we show that clusters of codons with biased codon usage patterns may serve as nucleic-acid-level domains that affect gene functions, just as a sequence of amino acids with a specific order and/or specific biochemical properties can form a protein domain. We accomplished this by developing a conservative statistical approach to identify PFCCs in the *D. melanogaster* genome. We have identified over 3000 PFCCs, and most of them strongly prefer rare codons. Nevertheless, we also found that a small proportion of the PFCCs exhibit other patterns of codon usage, such as preference for common codons, which was not reported before. We showed that although the PFCCs are associated with specific protein functional classes including transmembrane proteins and transcription factors, most of them are not associated with known protein domains defined by amino acid sequences. As a proof-of-principle, we used the example of a rare-codon cluster associated with the inactivation gate of Nav to propose a hypothesis concerning how a PFCC could affect specific biochemical and physiological properties of a protein. Together, our results suggest that it is likely a general phenomenon that codon clusters with biased codon usage patterns serve as diverse “hidden domains” involved in regulating protein functions.

In this paper, based on a widely used codon usage index CAI [15], we proposed an alternative codon usage index TCAI (see Materials and Methods: Calculating TCAI) that was used for classifying PFCCs. Compared to CAI, TCAI is better at describing the preference for rare codons. This is because when CAI is calculated, codon usage frequencies are all normalized to the frequencies of the most common synonymous codons. Thus, the CAI value of any codon cluster that strictly uses common codons will always be 1, while if two codon clusters that strictly use rare codons but have different amino acid sequences, they may have fairly different CAI values. However, by using the newly proposed TCAI, rare-codon clusters will have similar TCAI values that are −1 or very close to −1, while common-codon clusters keep TCAI values at 1 or near 1. Thus, TCAI is a good choice when researchers intend to identify rare-codon clusters.

In comparison to previous methods for detecting functional codon clusters [7], the method presented here is more conservative in terms of detecting rare-codon clusters due to the usage of both whole-genome and gene-specific codon usage patterns as the background codon usage. Yet, it is more powerful in terms of detecting other types of codon clusters due to a more relaxed assumption about the possible functional roles of codon clusters. The diverse codon usage patterns and locations of the PFCCs suggest that codon clusters may affect protein functions through various mechanisms. The major mechanism through which codon clusters regulate protein functions is possibly the deceleration of translation, as shown by the preponderance of rare-codon clusters in the identified PFCCs. However, we must admit that the preponderance of rare-codon clusters may be partly an artifact of technically easier detection of the preference for rare codons by our approach. To increase the power of codon-cluster-detection algorithms and more accurately assess the prevalence and importance of different types of codon clusters, researchers may need to incorporate phylogenetic analyses of homologous protein-coding genes in order to identify codon clusters with conserved codon usage patterns.

Consistent with previous reports [7], we found that some of the PFCCs are associated with known protein domains defined by amino acid sequences, which suggests that some codon clusters do have the potential to assist correct folding and modifications of protein domains. However, we also found that the majority of PFCCs are not associated with known protein domains [11,23], indicating that these PFCCs may carry necessary information for regulating protein functions and such information cannot be predicted from amino acid sequences. Thus, codon clusters could serve as “hidden domains” in protein-coding sequences. For example, some “free coiled regions” of proteins may not be actually “free”: their folding and modifications could be restricted by the codon usage patterns of the corresponding genomic regions. Further investigation into the codon clusters that may encode “hidden domains” could be important for biologists to better understand how genetic information directs the functions of proteins.

As we have shown by the example of rare-codon clusters in the DIII-IV linkers of Nav proteins, functional codon clusters may be important for some key functions of proteins. This could have important implications for molecular evolutionary studies and biological engineering practice. For molecular evolutionary studies, codon clusters with critical functions suggest that synonymous sites in such functional codon clusters may bias the estimation of the rate of neutral evolution if researchers consider synonymous mutations as neutral mutations. Moreover, it is possible that the selective pressure on synonymous codon usage may be even stronger than that on nonsynonymous mutations, which could greatly interfere the results and inferences of the evolutionary analyses based on the comparison between synonymous and nonsynonymous sites. For biological engineering practice, functional codon clusters suggest that when transgenes are designed, simple codon optimization [37], which generally uses common codons to encode amino acid residues, may not be the best choice to achieve desired structure and functions of the engineered proteins. Instead, the codon usage of different regions within a transgene may need to be more delicately controlled.

Together, our data support the broad existence of diverse and functional codon clusters that may affect protein functions and associated phenotypes through various mechanisms. In this regard, we suggest that functional codon clusters should be seriously considered if researchers are to thoroughly understand how genetic information is interpreted into functional, phenotypic, and evolutionary outputs *in vivo*.

## 5. Materials and Methods

### 5.1. Reference protein-coding sequences

Reference protein-coding sequences of *D. melanogaster* were downloaded from Ensembl 89 [38]. Protein-coding sequences fulfilling the following criteria were chosen. 1) The sequence length is a multiple of three. 2) The sequence uses standard genetic code. 3) For each gene, only the longest mRNA isoform was used; if there were multiple isoforms of the same length, then the first record shown in the FASTA file was used.

The protein-coding sequences of Nav1 and Nav2 in analyzed species can be found in Table S5.

### 5.2. Identifying PFCCs

Fig. S1 depicts how to identify PFCCs in a protein-coding sequence. For a window *W*_*i*_ starting with the *i*th codon in a protein-coding sequence, the window size *S* is set to vary between 5 to 50 codons. For each window size, two *χ*^2^ tests are performed by comparing the codon usage of the window respectively to whole-genome codon usage and gene-specific codon usage, and the higher *p*-value is selected as the representative *p*-value. Then the representative *p*-values are plotted against window sizes, which generates a *p-S* curve representing a function *p*(*S*) that describes the relationship between *p*-value and window size (Fig. S1A-D). If *p*(*S*) is monotonic, the lowest *p*-value together with its corresponding *S* are selected as the representative *p* and *S* for *W*_*i*_, namely *p*_*i*_ and *S*_*i*_; otherwise the *p*-value and the *S* that correspond to the lowest stationary point of *p*(*S*) are selected as *p*_*i*_ and *S*_*i*_. For the focal protein-coding sequence, all *p*_*i*_’s are corrected by setting the false discovery rate (FDR) [39] to 0.05 so as to get the corrected *p*-values *p*_*i,corrected*_’s; then windows with *p*_*i,corrected*_ values lower than the threshold 0.05 are detected as positive segments with unexpected codon usage patterns (Fig. S1E). Finally, isolated positive segments, together with the codon clusters generated by merging overlapped positive segments, are detected as PFCCs.

### 5.3. Calculating TCAI

To calculate the TCAI of a given sequence of codons, the background relative codon usage frequencies need to be calculated first. For example, if a gene uses 10 AAA and 30 AAG to encode Lys, the gene-specific background relative codon usage frequencies of AAA and AAG will respectively be 10/(10+30)=0.25 and 10/(10+30)=0.75. Then the focal sequence of codons is translated to an amino acid sequence. The next step is to generate a pseudo-sequence of codons according to the amino acid sequence and the background relative codon usage frequencies. For example, assuming that the amino acid sequence is Lys-Lys and the background relative codon usage frequencies are 0.25 for AAA and 0.75 for AAG, the first Lys will have a 25% chance to be encoded by AAA and 75% chance to be encoded by AAG, and so will the second Lys. This step of pseudo-sequence generation is repeated for 10,000 times so that there will be 10,000 pseudo-sequence of codons, which represent the expected results if codons are used randomly to encode the amino acids. Then the CAIs [15] of all pseudo-sequences and the CAI of the actual codon sequence are calculated. Finally, TCAI is calculated by subtracting the proportion of pseudo-sequence whose CAIs are higher than the CAI of the actual sequence from the proportion of pseudo-sequences whose CAIs are lower than the CAI of the actual sequence.

When TCAI is −1, it means that none of the pseudo-sequences has a CAI lower than the actual sequence; thus, the actual sequence strongly prefers rare codons. In contrast, when TCAI is 1, the actual sequence strongly prefers common codons.

### 5.4. K-mean clustering of PFCCs

K-mean clustering is done by using the online tool at http://scistatcalc.blogspot.com/2014/01/k-means-clustering-calculator.html. The number of clusters (i.e., K) is determined by the elbow method, according to https://pythonprogramminglanguage.com/kmeans-elbow-method/. Each input data point of K-mean clustering is specified by its gene-specific and whole-genome TCAIs.

### 5.5. Calculating RLI

For a protein-coding sequence with *L* codons, the RLI of a PFCC which starts at the *i*th codon and has a size of *S*_*i*_ codons is calculated as (*i* + *S*_*i*_ / 2) / *L*.

### 5.6. Searching for transmembrane helices

For a focal PFCC, the protein sequence from the first residue or the 150th residue upstream to the PFCC-encoded region, whichever is closer to the PFCC-encoded region, to the last sense codon or the 150th codon downstream to the PFCC-encoded region, whichever is closer to the PFCC-encoded region, is input to TMHMM [21] in order to search for transmembrane helices near or overlapped with the PFCC-encoded region. The coordinates of identified transmembrane helices are recorded.

### 5.7. Searching for Pfam protein domains

For a focal PFCC, the protein sequence from the first residue or the 150th residue upstream to the PFCC-encoded region, whichever is closer to the PFCC-encoded region, to the last sense codon or the 150th codon downstream to the PFCC-encoded region, whichever is closer to the PFCC-encoded region, is input to the hmmscan program of HMMER [23] on https://www.ebi.ac.uk/Tools/hmmer/search/hmmscan in order to search for Pfam protein domains [11] near or overlapped with the PFCC-encoded region. The coordinates and names of identified Pfam domains are recorded.

### 5.8. Classifying association between PFCCs and protein domains

The association between a PFCC and a protein domain is classified to one of the following categories.

1. No association: The closest distance between the PFCC-encoded region and the protein domain is longer than 20 residues.
2. 1-to-multiple association: Multiple protein domains are overlapped with the region that starts from the 20th residue upstream to the PFCC-encoded region and ends at the 20th residue downstream to the PFCC-encoded region.
3. Cluster in domain: Only one protein domain is associated with the PFCC. The PFCC-encoded region locates within the protein domain.
4. Domain in cluster: Only one protein domain is associated with the PFCC. The protein domain locates within the PFCC-encoded region.
5. Domain overlap left of cluster: Only one protein domain is associated with the PFCC. The start of the protein domain is upstream to the PFCC-encoded region and the end of the protein domain locates within the PFCC-encoded region.
6. Domain overlap right of cluster: Only one protein domain is associated with the PFCC. The start of the protein domain locates within the PFCC-encoded region and the end of the protein domain is downstream to the PFCC-encoded region.
7. Domain upstream to cluster: Only one protein domain is associated with the PFCC. The end of the protein domain is upstream to the PFCC-encoded region.
8. Domain downstream to cluster: Only one protein domain is associated with the PFCC. The start of the protein domain is downstream to the PFCC-encoded region.

### 5.9. Alignment of Nav orthologs and identification of DIII-IV linkers

Nav orthologs were aligned by using MAFFT algorithm [40,41]. The annotated DIII-IV linkers of Dmel/Nav1 (https://www.uniprot.org/uniprot/P35500) and Dmel/Nav2 (https://www.uniprot.org/uniprot/Q9W0Y8) were used to locate the DIII-IV linkers of the Nav1 and Nav2 in other analyzed species.

## 6. Acknowledgements

**General:** We thank Dr. Barak Cohen, Dr. Ian Duncan, Dr. David Queller, and Dr. Hani Zaher for helpful discussion. **Funding:** This work was supported by the National Institutes of Health (grant numbers R21NS089834 to Y.B.) and the National Science Foundation (grant numbers 1545778 and 1707221 to Y.B.). Z.P. is supported by the National Institutes of Health training program (grant number T32HG000045). **Author contributions:** Z.P. and Y.B. conceived the study; Z.P. performed the studies under the guidance of Y.B.; Z.P. and Y.B. wrote the manuscript and approved its final version. **Competing interests:** The authors declare no competing interests. **Data and materials availability:** computer code and raw data are available from Zhen Peng upon reasonable request.

## 10. Supplementary Material List

**Fig. S1. Using sliding window with adaptive size to identify PFCCs.**

**Table S1. Identified PFCCs.**

**Table S2. Analysis of the association between genes carrying N-terminal PFCCs (RLI<0.1) and GO terms.**

**Table S3. Analysis of the association between genes carrying PFCCs and GO terms.**

**Table S4. Analysis of the association between PFCCs and Pfam domains.**

**Table S5. Information of Nav genes.**

