## Supplementary figures and images for "Codon clusters with biased synonymous codon usage represent hidden functional domains in protein-coding DNA sequences"

### Fig_S1.png

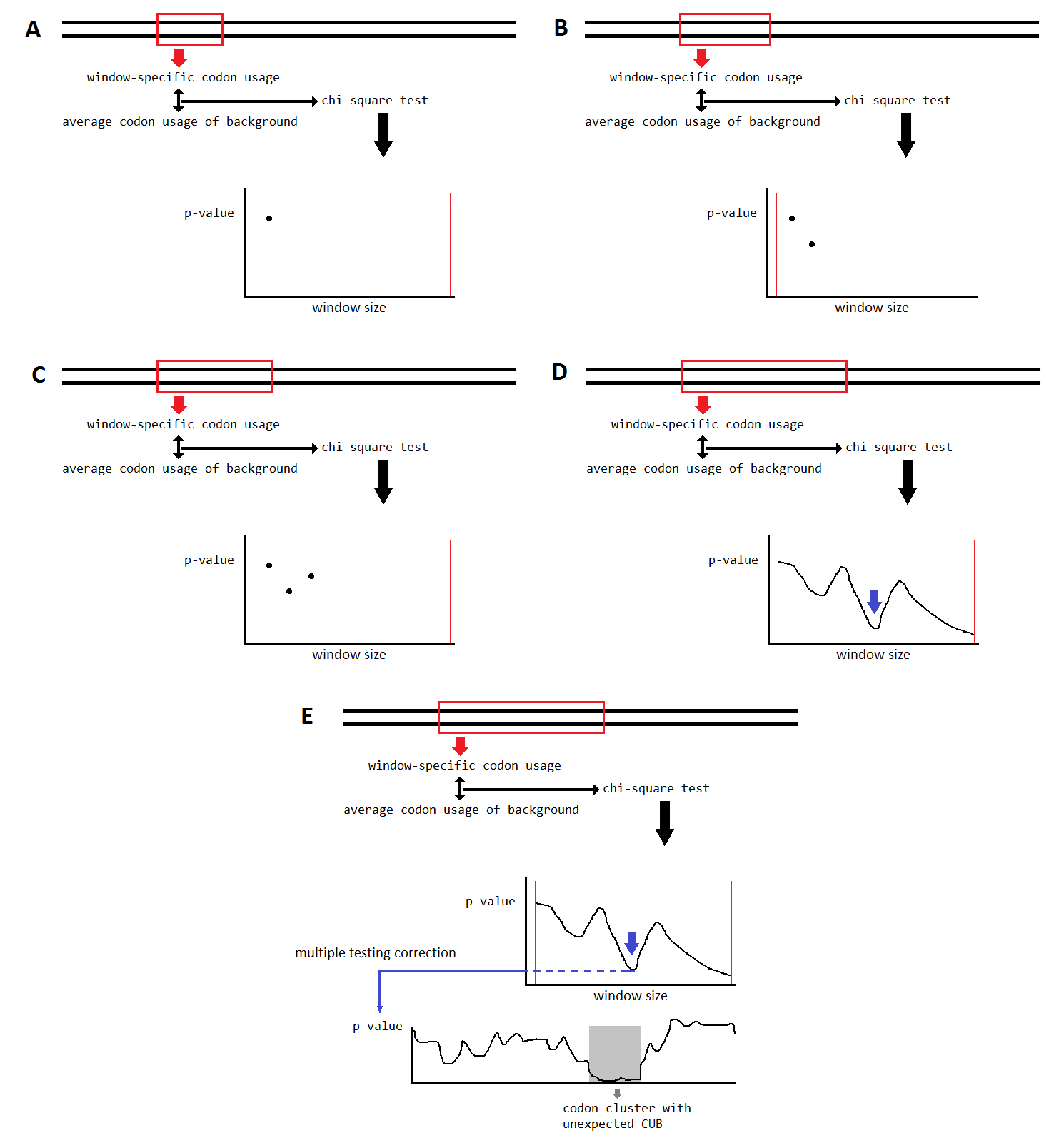
